# Template-independent genome editing and repairing correct frameshift disease *in vivo*

**DOI:** 10.1101/2020.11.13.381160

**Authors:** Lian Liu, Kuan Li, Linzhi Zou, Hanqing Hou, Qun Hu, Shuang Liu, Shufeng Wang, Yangzhen Wang, Jie Li, Chenmeng Song, Jiaofeng Chen, Changri Li, Haibo Du, Jun-Liszt Li, Fangyi Chen, Zhigang Xu, Wenzhi Sun, Qianwen Sun, Wei Xiong

## Abstract

Frameshift mutation caused by small insertions/deletions (indels) often generate truncated and non-functional proteins, which underlies 22% inherited Mendelian disorders in humans. However, there is no efficient *in vivo* gene therapy strategies available to date, especially in postmitotic systems. Here, we leveraged the non-homologous end joining (NHEJ) mediated non-random editing profiles to compensate the frameshift mutation in a USH1F mouse model – *av3j*. After treatment by the selected gRNA, about 50% editing products showed reading-frame restoration, and more than 70% targeted hair cells recovered mechanotransduction. *In vivo* treatment ameliorated the hearing and balance symptoms in homozygous mutant mice. Furthermore, a scale-up analysis of 114 gRNAs targeting 40 frameshift deafness mutations reveals that 65% loci have at least one gRNA with predicted therapeutic potential. Together, our study demonstrates that the NHEJ-mediated frame restoration is a simple and highly efficient therapeutic strategy for small-indel induced frameshift mutations.

## Main

The CRISPR/Cas9 based genome-editing techniques enable engineering pathogenic DNA variants, which has drawn significant attention on the application for gene therapy in live animals^1,2^. The base-editing techniques achieve correction of substitution mutations^3–5^. Homology-dependent strategies can precisely correct mutations according to the given templates^6,7^. And homology-independent editing strategies can introduce large insertion in targeted region, thus leading to ameliorate disorders caused by large deletions^8^. However, specific and efficient therapeutic strategies for frameshift mutations are still lacking. Following CRISPR/Cas9 cleavage, the induced DNA double strand breaks (DSBs) are repaired by two major endogenous DNA repair pathways, precise homology-directed repair (HDR) and error-prone non-homologous end-joining pathway (NHEJ)^2^. Although the HDR pathway can restore any mutations based on the given donor templates, its application is largely limited due to low recombination efficiency and is almost absent in non-dividing, terminally differentiated cells/organs^2^. While NHEJ is the major repair mechanism but introduces stochastic indels, thus it has been widely applied in reading frame disruptions. However, emerging evidence in dividing cells has shown that the editing profiles through end-joining pathways are non-random and mainly gRNA-sequence dependent^9–14^. Furthermore, gRNAs containing dominant indels could be predicted by machine learning models^10,12,13^, which can compensate the pathogenic frameshift mutations, as exhibited in patient-derived fibroblasts^12,15^.

Here we adopted this NHEJ-mediated frame-restoration strategy and evaluated its therapeutic potential *in vivo* within the context of hearing disorders. About 300 million people suffer from hearing loss worldwide, and 50% prelingual hearing loss are caused by genetic mutations^16^. Among them, Usher Syndrome is one of the most devastating diseases, which affect not only hearing, but also vision and balance function^17^. However, there is no pharmacological treatment available to date, especially for those types involving mutations in large proteins (*USH1B*, MYO7A, 254 KD; *USH1D*, CDH23, 370 KD; *USH1F*, PCDH15, 216 KD; *USH2C*, VLGR1, 693 KD)^17^. In order to explore the frameshift correction ability of this strategy for huge protein in postmitotic system, we recruited an autosomal recessive USH1F mouse model – *av3j*, in which a spontaneous A insertion causes frameshift in the 7.9 kb *Pcdh15* transcript and results in truncated PCDH15 protein lacking transmembrane and intracellular domains^18^ (Fig. 1a). PCDH15 is one of the two components forming the tip link that gates the mechanotransduction channels in inner-ear hair cells^19^, including cochlear outer hair cells (OHCs), inner hair cells (IHCs) and vestibular hair cells (VHCs). The *av3j* mutant hair cells cannot detect mechanical force thus the homozygous mice show profound congenital deafness and severe circling behavior.

**Fig. 1.**
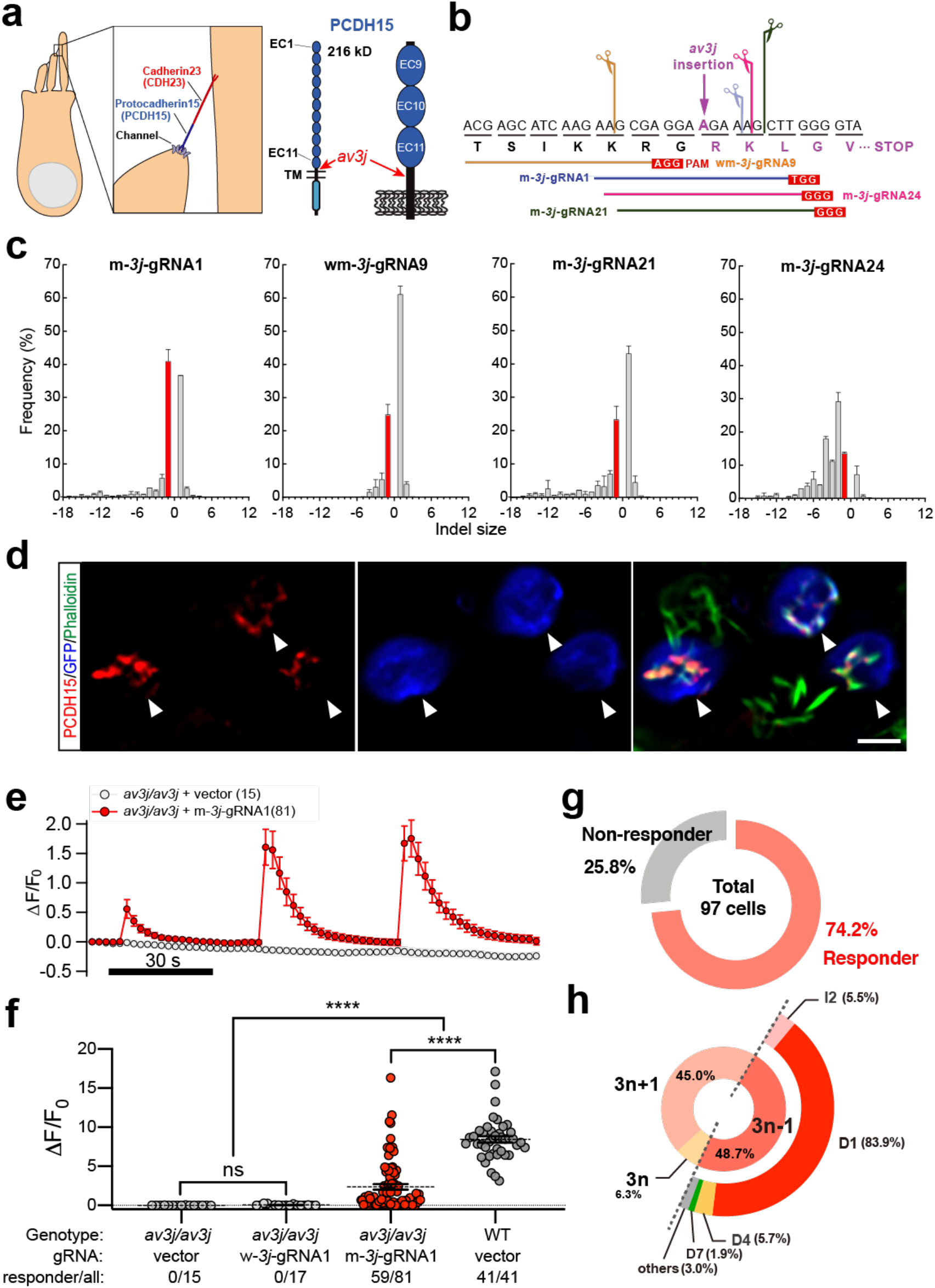
The selected m-*3j*-gRNA1 efficiently restores the *av3j* deficiency *ex vivo*. **a,** PCDH15 is one of the two components of the tip link that gates the mechanotransduction channel in hair cells. The 216-kD PCDH15 contains 11 extracellular cadherin (EC) repeats and a transmembrane (TM) domain. The *av3j* mutation brings an A insertion between the EC11 and TM domain and causes an early stop codon in the *Pcdh15* gene. **b,** Four 17-nt gRNAs were designed to target the *av3j* A insertion site. The m-*3j*-gRNA1, m-*3j*-gRNA21, and m-*3j*-gRNA24 cover the insertion, while wm-*3j*-gRNA9 can target both wild-type (WT) and *av3j* alleles. **c,** The editing profiles of the 4 gRNAs shown as the indel frequency at each indel size. The bars of 1-bp deletion were highlighted in red, which was 40.9% for m-*3j*-gRNA1. Each gRNA profile had 2-3 replicates. Error bars, SD. **d,** PCDH15 protein immunostaining in the electroporated P3 *av3j/av3j* cochleae cultured for 2 days *in vitro* (P3+2DIV). PX330 containing m-*3j*-gRNA1 and Cas9 (PX330-m-*3j*-gRNA1) was co-electroporated with N1-EGFP. Note that only the hair bundles of electroporated *av3j/av3j* hair cells showed obvious signals (arrowhead). Scale bar, 3 μm. **e,** Averaged traces of Ca^2+^ responses activated by fluid-jet stimuli in *av3j/av3j* OHCs electroporated by control PX330 vector or PX330-m-*3j-*gRNA1 (P1+4 DIV). Error bars, SEM. **f,** Quantification of the maximal Ca^2+^ responses in OHCs from recordings similar to (**e**). W-*3j*-gRNA1 targets the corresponding WT sequence and shares the same PAM with m-*3j*-gRNA1. Responsive and total recorded cell numbers are shown in panels. Brown-Forsythe and Welch ANOVA test; ns, no significance, *\*\*\*\*P* < 0.0001, error bars, SEM. **g,** 74.2% (72/97) *av3j/av3j* auditory hair cells, including both OHCs (**f**) and IHCs (Supplementary Figure 1c), recovered mechanosensitivity after PX330-m-*3j*-gRNA1 electroporation. **h,** 48.7% editing products in m-*3j-*gRNA1 are ‘3n-1’, which could restore the reading frame of *av3j*, and were dominated by 1-bp deletion (D1), 4-bp deletion (D4) and 2-bp insertion (I2) (See Supplementary Fig. 1d for details). Note that, correction of only one allele is sufficient to achieve a functional recovery of the diploid *av3j/av3j* hair cells.

By gRNA screening in cochlear explants, we identified one gRNA with half of its profiles could compensate the 1-bp insertion in *av3j* mice. After genome editing, the PCDH15 protein expression and localization were restored, and nearly 3/4 targeted hair cells recovered mechanotransduction. Early postnatal delivery of the gRNA-containing virus in scala media successfully ameliorate auditory and vestibular symptoms of *av3j* homozygous mice. A scale-up gRNA screening on postmitotic cochlear tissues revealed that this frame-restoration strategy is suitable for more than 60% of the tested frameshift deafness sites. Our results demonstrate that the NHEJ-mediated frame restoration is a simple and highly efficient approach to correct small-indel induced frameshift mutations, which opens up new perspectives for gene therapy strategy targeting frameshift mutations, especially in postmitotic systems.

## Result

### Design and characterization of gRNAs for frameshift restoration *ex vivo*

By searching the NGG PAM sequence proximal to the inserted A, four gRNAs targeting *av3j* mutation were designed (Fig. 1b). The gRNAs with 17 nucleotides were used for higher editing specificity^20^. In order to interrogate the genome editing outcomes in inner-ear cells, we established an *ex vivo* gRNA screening system which combined cochlea culture, injectoporation^21^, FACS, and amplicon sequencing (Supplementary Fig. 1a). All designed gRNAs show unique editing profiles, in which m-*3j*-gRNA1, wm-*3j*-gRNA9, and m-*3j*-gRNA21 preferred one or two dominant indel sizes while m-*3j*-gRNA24 had multiple infrequent outcomes (Fig. 1c). Distinct profiles can be found between the gRNAs with even 1-bp shift (m-*3j*-gRNA1, m-*3j*-gRNA21, and m-*3j*-gRNA24). Considering *av3j* is a 1-bp insertion, we selected m-*3j*-gRNA1 that exhibits the highest percentage of 1-bp deletion (D1) (red bars, Fig. 1c).

Next, we tested whether m-*3j*-gRNA1 could mediate functional restoration in mutant *av3j/av3j* hair cells. Take advantage of our tissue injectoporation system, we electroporated m-*3j*-gRNA1 and wildtype spCas9 (PX330-m-*3j*-gRNA1) in postnatal day 3 *av3j/av3j* cochleae and cultured for 2 days *in vitro* (P3+2DIV). PCDH15 puncta was specifically detected in some electroporated *av3j/av3j* OHCs, at their hair bundle tips, but not in the surrounding unelectroporated hair cells (Fig. 1d). GCaMP-based Ca^2+^ imaging was further applied to assess the mechanotransduction recovery of electroporated *av3j/av3j* hair cells. Surprisingly, among the 97 tested cochlear hair cells, including 81 OHCs (Fig. 1e,f) and 16 IHCs (Supplementary Fig. 1c), 72 restored responses to the mechanical stimulation (74.2%, Fig. 1g), which was much higher than the 40.9% D1 events showed in editing profiles (Fig. 1c), indicating that other frame-restored products might be functional too.

### Function evaluation of the top frame-restored editing products for m-*3j*-gRNA1

To investigate the function of other frame-restored editing products, we further analyzed the editing products of m-*3j*-gRNA1. There are mainly three frame-restored types (refers to products with the same indel sizes): D1, 4-bp deletion (D4) and 2-bp insertion (I2), and each frame-restored types contain 1-2 top products (Supplementary Fig. 1d). We constructed the top editing products of the three frame-restored types: PCDH15-E1373R (D1-top1), PCDH15-E1373del (D4-top1) and PCDH15-E1373RK (I2-top1) (Supplementary Fig. 1e), and further examined the functional recovery by electroporation and Ca^2+^ imaging on *av3j/av3j* hair cells (Supplementary Fig. 1f,g). All frame-restored constructs rescued the mechanosensitivity of electroporated *av3j/av3j* hair cells, though the Ca^2+^ responses were smaller in PCDH15-E1373RK (I2-top1) comparing to the wildtype PCDH15 control (Supplementary Fig. 1e-g). These results show that almost all top frame-restored editing products of m-*3j*-gRNA1 can mediate functional rescue of *av3j/av3j* hair cells, though containing 1-2 amino acid alterations compared with WT protein sequences. All the frame-restored products (3n-1) occupied 48.7% of total indels in m-*3j*-gRNA1 (Fig. 1h). For mammalian cells are diploid and *av3j* heterozygous hair cells are fully functional, so the function restoration rate of targeted hair cells would be 48.7%*48.7%+2*48.7%*(1-48.7%)=73.7%, which is consistent with the 74.2% Ca^2+^ response recovery rate in the electroporated hair cells. These results reveal that our frame-restoration strategy enables highly efficient reading-frame correction and function restoration in postmitotic cells.

### *In vivo* editing profiles of m-*3j*-gRNA1 by virus delivery

To achieve the genome editing *in vivo*, we packed the U6-gRNA scaffold into the AAV2/9 capsid, with a mCherry expression cassette included to indicate virus transfection. Firstly, a preliminary transfection was applied to WT mice by scala media injection at P0-P2. The mCherry expression was observed in 100% IHCs in all observed cochleae at 14 day-post-delivery (PVD), and the transfection rates in OHCs and VHCs was 70% and 60%, respectively (Supplementary Fig. 2).

Then we assessed the *in vivo* genome editing profiles of m-*3j*-gRNA1 in *av3j/av3j* mice with *Cas9* knockin background^22^ (*av3j/av3j;Cas9+*). The cochlear and vestibular tissues were harvested at 2, 4, 8, 12, and 16 weeks PVD (Supplementary Fig. 3a-d). The averaged editing efficiencies were 60% for cochlear tissues and 70% for vestibular organs, which reached a plateau as early as 4 weeks, with the major editing events carried out within 2 weeks (Fig. 2b). When comparing the detailed editing profiles in cochleae and vestibules *in vivo* with that from cochleae *ex vivo*, we found that D1 and I1 were dominant at all tested conditions. Although the exact frequencies vary (Fig. 2c-e), the dominant frame-restored products are exactly the same, as manifested by the indel positions and inserted bases (Fig. 2f-h). Besides, the editing outcomes are more similar in profiles from cochleae (*in vivo* and *ex vivo*) compared to those from vestibular organs, indicating a system-specific bias in editing profiles.

**Fig. 2.**
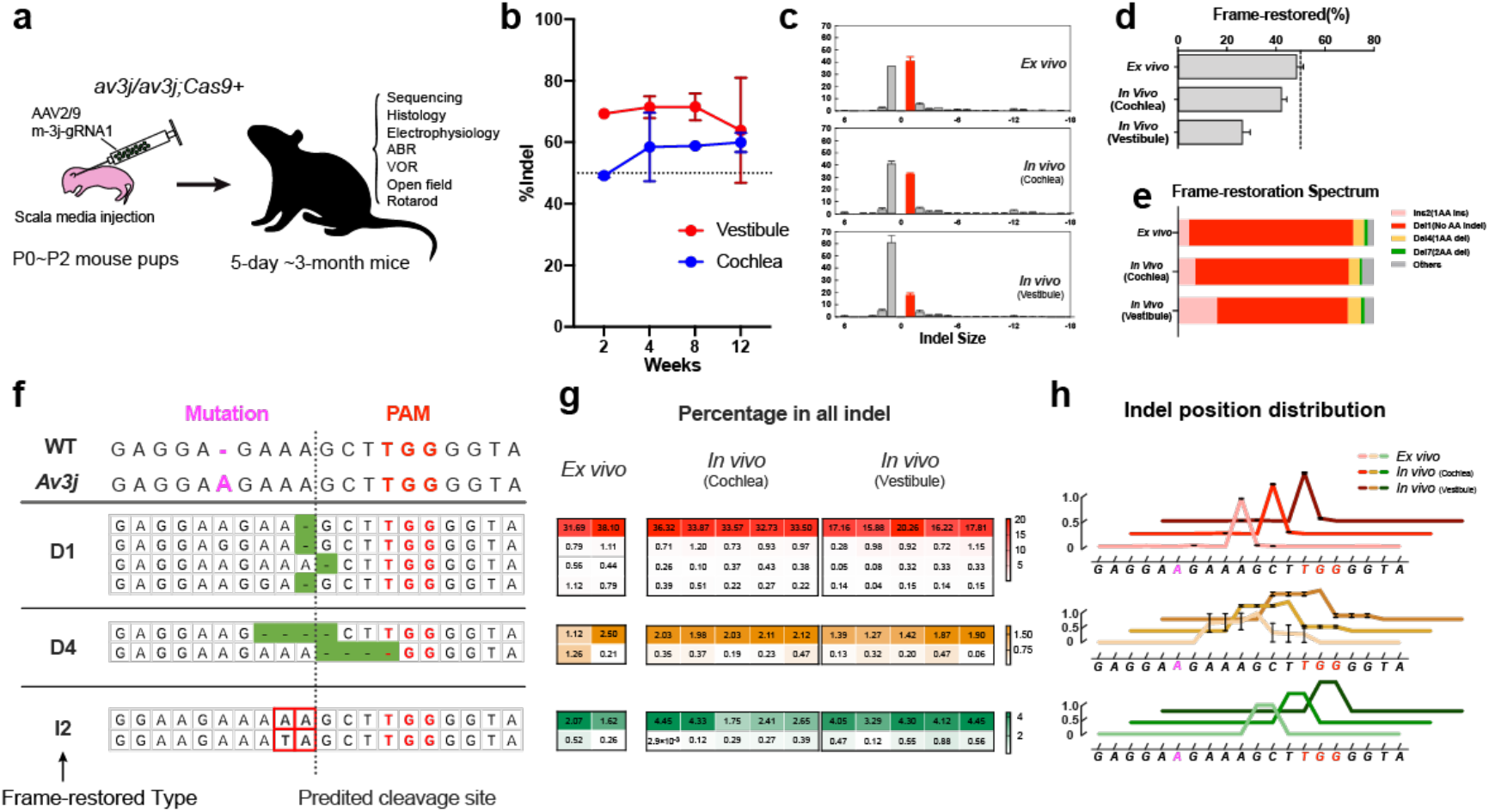
Similar editing outcomes of m-*3j*-gRNA1 *ex vivo* and *in vivo*. **a,** A cartoon illustrating the overall procedures of *in vivo* gene delivery and evaluation in Fig. 2-4. AAV2/9-U6-m-*3j*-gRNA1-CMV-mCherry (0.8 μl, 1.5 × 10^14^ vg/ml) was injected to scala media of P0-P2 *av3j/av3j;Cas9+* mice bilaterally, followed by a variety of measurements at indicated ages. **b,** Genome editing efficiencies (quantified as indel percentage) in virus transfected cells of cochleae and vestibules. The tissues were harvested at age of 2, 4, 8, 12, 16 weeks post-viral delivery (PVD). The transfected cells were sorted by FACS then followed by deep sequencing (Supplementary Fig. 3a-d for details). Each group has 2-5 animals. **c-h,** Detailed editing product analysis of m-*3j*-g1 in electroporated cochlear cultures (P1+4DIV, *ex vivo*), AAV transfected cochleae and vestibules from *av3j/av3j;Cas9+* mice (1 month PVD, *in vivo*). Error bars, SD. **c,** The editing profiles of m-*3j*-gRNA1. Error bars, SD. **d,** The percentage of frame-restored editing product of m-*3j*-gRNA1. **e,** Frame-restoration spectrum analysis. **f,** The top three frame-restored types were dominant by 1-2 editing products. The mutant and WT sequence are shown on the top. *Av3j* mutation and PAM sequence are illustrated in magenta and red. The predicted Cas9 cleavage site is indicated by a dash line at 3bp upstream the NGG PAM. **g,** The percentage of the corresponding editing products in all indels, with each column indicating a single replicate. **h,** The indel position analysis. Color: lightest, cochleae *ex vivo*; medium, cochleae *in vivo*; darkest: vestibules *in vivo*. Error bars, SD.

As previous study showed a recut of small indel products to larger deletions^9^, we studied the editing profiles along time, and found that the outcomes of virally-delivered m-*3j*-gRNA1 were stable at all tested time points (Supplementary Fig. 3e,f). We also tested the off-target effects for this long-term AAV-mediated expression by amplicon sequencing of predicted off-target sites (see Methods). 2 out of 16 selected off-target sites show detected editing with averaged indel frequencies at 0.7% and 0.3% (Supplementary Fig. 3g,h). These results demonstrate that our *ex vivo* screening and evaluation indeed help assess the therapeutic potential of gRNAs for the frame-restoration gene therapy *in vivo*.

### NHEJ-mediated frame restoration recovers mechanotransduction of *av3j* hair cells *in vivo*

We initially examined the PCDH15 protein expression and localization in treated mice after virally delivering m-*3j*-gRNA1 at P0-2. The PCDH15 protein expression is recovered after the treatment, as PCDH15 puncta was observed in the hair bundle tips of the transfected *av3j/av3j;Cas9+* OHCs at 5 days PVD (Fig. 3a).

**Fig. 3.**
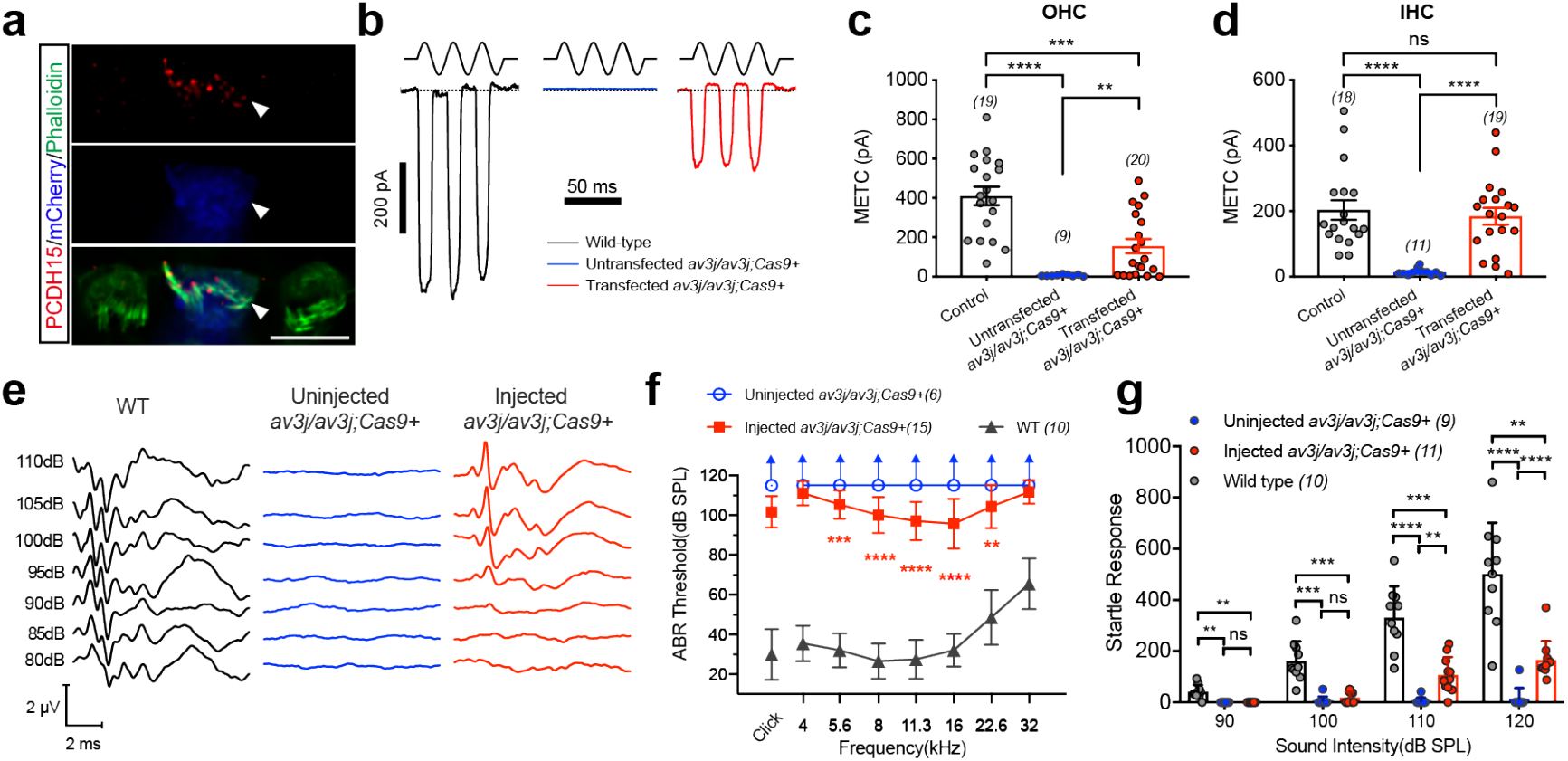
*In vivo* delivery of m-*3j*-gRNA1 ameliorates auditory symptom of *av3j/av3j;Cas9+* mice. **a,** PCDH15 immunostaining (red) showing recovery of protein expression in the stereocilium tips in transfected OHCs (blue) from P0-P2 AAV injected *av3j/av3j;Cas9+* mice (tissue harvested 5 days PVD). Hair bundle were stained with phalloidin (green). Note that only the hair bundles of transfected *av3j/av3j* hair cells showed obvious signals (arrowhead). Scale bar, 6 μm. **b,** Representative examples of mechanotransduction currents recorded from OHCs in control and *av3j/av3j;Cas9+* mice with or without transfection (10 days PVD). A fluid jet was used to stimulate the hair bundle during recordings. **c-d,** Quantification of the mechanotransduction current amplitudes from recordings similar to (**b**) in OHCs (**c**) and IHCs (**d**). 14 out of 20 OHCs and 18 out of 19 IHCs were responsive. Littermate *av3j/+;Cas9+* or WT mice at the same age were used for control. **e,** Representative click ABR responses from 4-5 week WT, uninjected and injected *av3j/av3j;Cas9+* mice. **f,** Mean ABR thresholds in WT, uninjected and injected *av3j/av3j;Cas9+* mice. Statistics represented the difference between uninjected and injected *av3j/av3j;Cas9+* group. 15 out of 33 injected *av3j/av3j;Cas9+* mice recovered ABR responses. Only mice with ABR responses were plotted in the injected group. Arrows indicate the thresholds are higher than the maximal stimulus level. **g,** Startle responses to white noises were recorded in 4-5 week WT, uninjected and injected *av3j/av3j;Cas9+* mice. Only mice with startle responses were plotted in the injected group. 17 injected mice were tested in startle responses prior to ABR test, in which 11 had both startle and ABR recovery, 4 with ABR recovery only, and the rest 2 with no restoration in both tests. Brown-Forsythe and Welch ANOVA test; error bars, SEM. ns, no significance, *\*P* < 0.05, *\*\*P* < 0.051, *\*\*\*P* < 0.001, *\*\*\*\*P* < 0.0001.

To investigate the recovery of mechanotransduction, fluid-jet evoked currents were monitored by whole-cell patch-clamp recording on auditory hair cells at 10 days PVD (Fig. 3b). Among these transfected hair cells, 14 out of 20 OHCs and 18 out of 19 IHCs restored mechanotransduction currents (Fig. 3c,d), which is consistent with the high recovery efficiency observed *ex vivo*.

To assess the *in vivo* functional rescue ratio in targeted auditory hair cells, we used styryl FM5-95 dye to label the function-restored hair cells. Styryl FM dye has been reported entering to hair cells through mechanotransduction channels, thus it is used as an indicator for channel opening^23^. In total of 929 counted transfected OHCs, 78.3% recovered dye uptake (Supplementary Fig. 4a,b). As previous study has shown a gradual hair cell loss in *av3j* homozygous mice^24^, to evaluate whether our treatment could reverse this symptom, we next quantified the hair-cell survival in injected cochleae. After treatment, hair-cell survival was improved in the apical turn, while the cell death in the middle and basal turn was not significantly reversed (Supplementary Fig. 4c-g).

### NHEJ-mediated frame restoration recovers auditory function of *av3j* mice *in vivo*

To evaluate whether the auditory function of injected *av3j/av3j;Cas9+* mice was rescued, we recorded the auditory brainstem response (ABR) at 4-5 week PVD. 15 out of 33 injected mice showed obvious ABR responses, compared to totally no signals in the uninjected animals (Fig. 3e-d). And the maximum click ABR threshold improvement was up to 20 dB, with 3 best mice harbored pure-tone ABR threshold at 80 dB SPL for 11.3 kHz. However, the lack rescue of the other 18 injected mice was unclear, may have been due to lower virus transfection rates.

In order to evaluate the recovery of whole auditory circuits, acoustic startle responses were tested by random white noise stimulation of multiple sound intensities. Obvious startle responses could be recorded in some injected mutant mice upon large stimuli. 17 injected *av3j/av3j;Cas9+* mice were tested in acoustic startle test prior to ABR recording, in which 11 had both startle and ABR recovery, 4 with ABR recovery only, and the rest 2 with no restoration in both assays (Fig. 3g). These data demonstrate that highly efficient, postnatal recovery of hair-cell mechanotransduction could reverse the totally abolished hearing loss in *av3j* mice.

### NHEJ-mediated frame restoration recovers vestibular function of *av3j* mice *in vivo*

As *av3j* mice also has severe balance problem due to totally abolished mechanotransduction in VHCs, we further evaluated the functional rescue in vestibular system. In the transfected *av3j/av3j;Cas9+* VHCs, protein expression of PCDH15 was also detected at 10 days PVD (Fig. 4a). In 11 transfected VHCs, 7 restored mechanotransduction currents (10 days PVD, Fig. 4b-c).

**Fig. 4.**
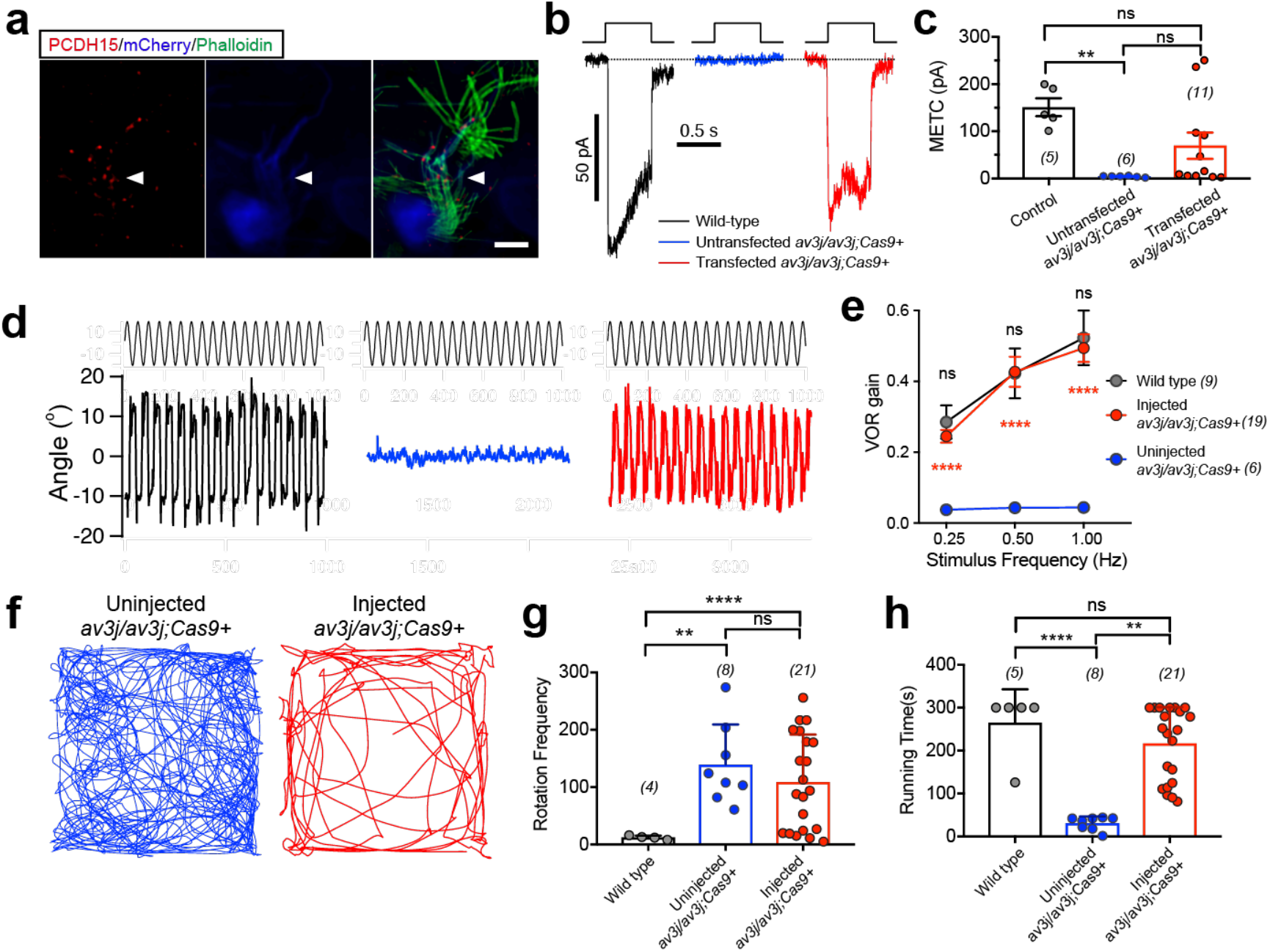
*In vivo* delivery of m-*3j*-gRNA1 ameliorates vestibular symptom of *av3j/av3j;Cas9+* mice. **a,** PCDH15 protein expression (red) was recovered in the stereocilium tips of transfected vestibular hair cells (VHCs, blue) from injected *av3j/av3j;Cas9+* mice (10 days PVD). Hair bundle stained with phalloidin (green). Note that only the hair bundles of transfected *av3j/av3j* hair cells showed obvious signals (arrowhead). Scale bar, 3 μm. **b,** Representative examples of mechanotransduction currents recorded from VHCs in WT mice, uninjected and injected *av3j/av3j;Cas9+* mice (10 days PVD). A fluid jet was used for stimulation. **c,** Quantification of the mechanotransduction current amplitudes from recordings similar to (**b**). For transfected *av3j/av3j;Cas9+* VHCs, 7 out of 11 (64%) were responsive. P10 WT were used for positive control. **d,** Representative traces of VOR responses from WT mice, uninjected and injected *av3j/av3j;Cas9+* mice. **e,** Quantification of the VOR gain. Note that the responses of injected animals show no difference with WT mice in all frequencies. Statistics result between WT and injected group are shown in grey, injected vs uninjected shown in red. **f,** Examples of open-field locomotion traces of uninjected *av3j/av3j;Cas9+* littermates and injected *av3j/av3j;Cas9+* mice. **g,** Quantification of circling frequencies during open field tests. 7 out of 21 injected mice had similar circling frequencies as WT, indicating a total recovery of the circling symptom. **h,** Quantification of the running time in rotarod test obtained in WT mice, uninjected and injected *av3j/av3j;Cas9+* mice. Brown-Forsythe and Welch ANOVA test; ns, no significance, *\*\*P* < 0.01, *\*\*\*P* < 0.001, *\*\*\*\*P* < 0.0001; error bars, SEM.

To evaluate the overall vestibular function in injected *av3j/av3j;Cas9+* mice, we recorded the vestibulo-ocular reflex (VOR), in which the vestibular system detects head motions and triggers opposite eye movements to help vision stabilization. The injected mutant mice showed VOR response recovery close to wild-type levels at all tested rotation frequencies, in contrast to the totally absent VOR in the uninjected mice (Fig. 4d-e). Then we performed open field and rotarod test to examine the restoration of balance behavior. The circling frequencies were also reduced in some injected *av3j/av3j;Cas9+* mice, as 7 out of 21 animals almost stopped circling (Fig. 4f-g). In rotarod test, all injected *av3j/av3j;Cas9+* animals stayed longer on the rotating rod comparing to the uninjected mutants (Fig. 4h). And 6 injected mice could still stand on the rod when the tests had finished at 300 s just like the majority of WT animals, indicating almost full recovery of the balance function.

### Therapeutic gRNA screen for other frameshift deafness mutations

After the successful therapeutic application of this frame-restoration strategy on *av3j* mice, we further explored whether it can be extended to target other similar disease mutations with our *ex vivo* gRNA screening system (Supplementary Fig.1a). A total of 114 gRNAs were designed to target 40 human pathogenic deafness mutations that are caused by 1-bp indels, and only the highly similar sequences between humans and mice were selected (Supplementary Table 1,2). During the gRNA screening for *av3j* site, we found that the indel distributions and indel positions were very similar between gRNA pairs targeting the same genomic sites in *av3j* genome and WT genome, which share the same PAM positions and differ only in the 1-bp A insertion (Supplementary Fig. 5a-f). We next used the published genome editing prediction models, inDelphi^12^ and FORECasT^10^, to predict the editing profiles of 114 gRNA pairs targeting mutant genomes and WT genomes. Most gRNA pairs showed high correlations of their editing profiles (median Pearson *r*: inDelphi, 0.84; FORECasT, 0.90, Supplementary Fig. 5 g-j). All these indicate that the mutant and WT gRNA pairs share similar editing profiles. As the *ex vivo* results resemble the *in vivo* editing profiles (Fig. 2), electroporation on WT cochlear tissues was applied to investigate the editing profiles of the designed gRNAs.

For frame-restoration gRNA selection, multiple factors were considered in defining the therapeutic potential (Fig. 5a). Firstly, only gRNAs that mediate successful genome editing have the possibility of frameshift correction (editing efficiency). And fraction of frame restoration provides a baseline for function-rescue evaluation. For those sites tolerant for small amino acid changes, this factor may directly reflect the function-rescue fraction, thus enabling a quick evaluation of available gRNAs. Then the function of frame-restored products, which lays the foundation of final therapeutic outcome, is the key point in gRNA evaluation.

**Fig. 5.**
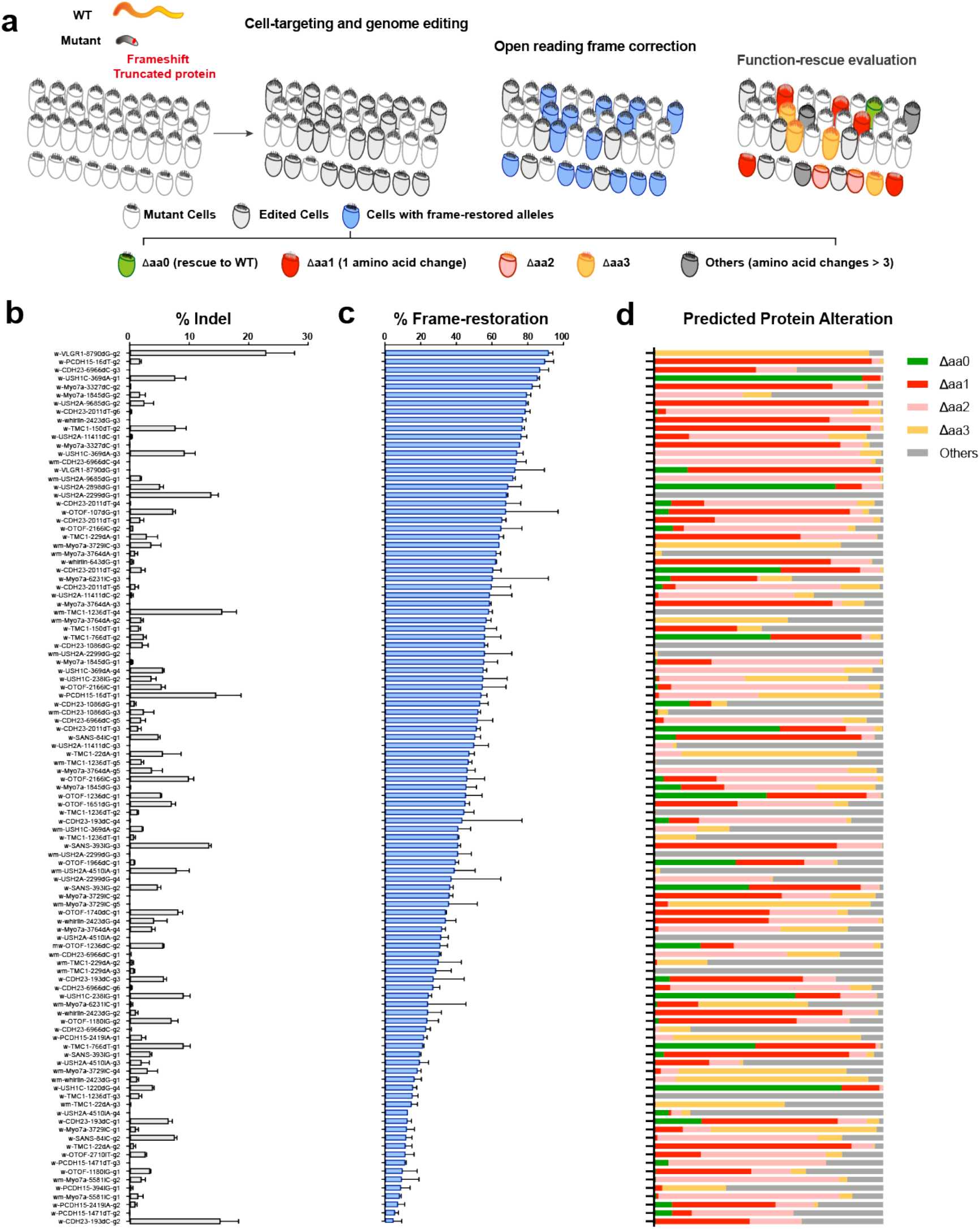
Editing outcome evaluation of 114 designed gRNAs targeting other frameshift deafness mutations. **a,** Cartoon showing multiple factors relevant for frame-restoration gRNA evaluation: genome editing efficiency, frame-restoration ratios, and editing product function-rescue evaluation. **b,** Genome editing efficiencies of designed gRNAs, in which the %Indel data come from N2a cell line 72h post transfection of gRNA-carrying PX330 plasmid for it was difficult to tightly control the percentage of electroporated tissue cells throughout FACS sorting. **c,** The percentage of predicted frame-restored products of designed gRNAs. **d,** The predicted protein alteration spectrums after genome editing of each gRNA based on deep sequencing data from electroporated tissues. Color code indicates the number of amino acid changes relative to WT.

All designed gRNA showed different editing efficiencies (Fig. 5b) and fraction of predicted frame restoration (Fig. 5c). As for the function of the frame-restored editing products, we quantified the predicted amino acid changes (Fig. 5d), and assumed that with fewer amino acid changes, the products have higher possibilities to achieve a final functional rescue. Besides the absolute altered numbers, the functional importance of affected amino acids is also relevant for functional evaluation of the editing products. So we did conservation analysis of each amino acid position near the mutant sites^25^. In order to link the conservation status and amino acid change number to the final protein function, we selected 24 editing products targeting 6 deafness sites, and validated their protein functions in either HEK293T cells or electroporated cochlear hair cells (Supplementary Fig. 6). In general, with lower conservation of altered amino acid, there is a higher possibility of protein function preservation. Some unconserved regions are tolerant for as many as 3 amino acid alterations, including both deletions and substitutions. And in many cases, deletions were more deleterious than substitutions (Supplementary Fig. 6). We incorporated all our observations into one functional prediction algorithm to help briefly evaluate the fraction of function-restored editing products in each gRNA (for details, see Methods).

### Radar plot assists evaluation and selection of therapeutic gRNAs for frameshift mutations

Finally, we integrated all these parameters into one radar plot to facilitate the therapeutic gRNA evaluation and selection (Fig. 6). The radar plot has 5 axis: %*Indel*; *%Frame-restoration*; *%Predicted aa changes* ≤3, showing percentage of editing products with equal or less than 3 amino acid changes compared with WT protein sequences; *%Predicted functional rescue*, the fraction of products with a predicted functional rescue; *Specificity*, a specificity score, which based on sequence homology across mouse genome, was calculated by published algorithms^26^ (for details, see Methods). From radar plot, w-*TMC1*-229dA-g1 was predicted as the best for *TMC1* c.229delA mutant site (Fig. 6a,d,g). For *PCDH15*-16dT, which locates in highly variable region, both two designed gRNAs could mediate high fractions of predicted function-rescue products (Fig. 6b,e,h). And for USH1C c.369delA mutation, w-*USH1C*-369dA-g1 would have the best performance (Fig. 6c,f,i). Note that, cleavage sites of w-*USH1C*-369dA-g3 and w-*USH1C*-369dA-g4 only separate 1-bp away, and they generate many identical editing products. Although w-*USH1C*-369dA-g3 has more frame-restored products and similar protein alteration spectrum compared with w-*USH1C*-369dA-g4, it contains the lowest functional rescue products, for w-*USH1C*-369dA-g3 preferred more amino acid deletions instead of substitutions compared with w-*USH1C*-369dA-g4 (Supplementary Fig. 6c,d).

**Fig. 6.**
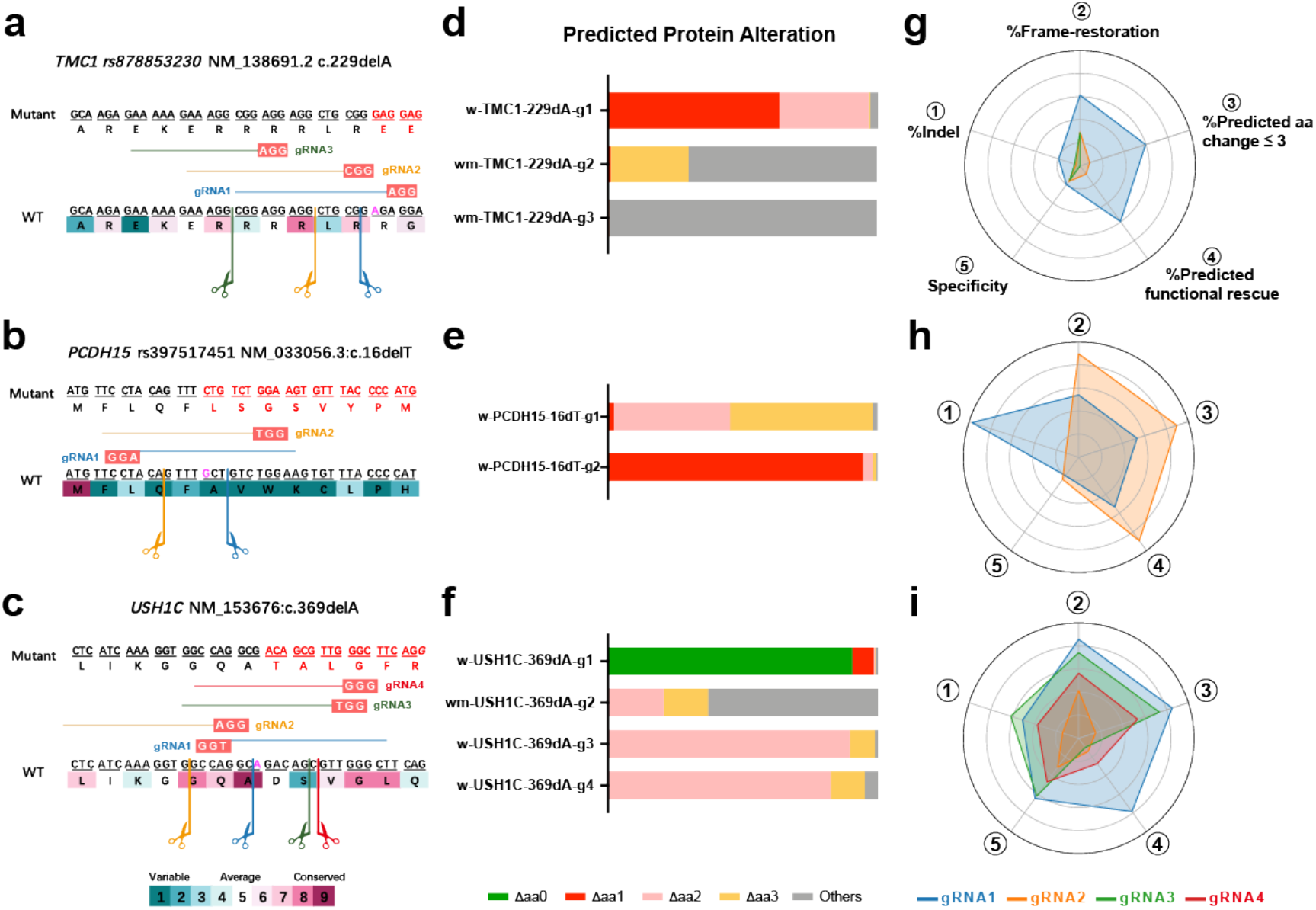
Evaluation and selection of therapeutic gRNAs targeting other frameshift deafness mutations. **a-c,** Therapeutic gRNA design for *TMC1* c.229delA (**a**), *PCDH15* c.16delT (**b**) and *USH1C* c.369delA (**c**). Magenta in WT indicate the mutant nucleotides, and frameshift sequences are colored in red in mutant sequences. The open reading frames are illustrated by underlines, and amino acids are colored by conservation levels, highly variable (deep green), highly conserved (muddy purple). The dash line showed the predicted cleavage site of each gRNA. **d-f**, The predicted protein alteration spectrum of gRNAs targeting *TMC1* c.229delA (**d**), *PCDH15* c.16delT (**e**) and *USH1C* c.369delA (**f**). **g-i,** Radar plots incorporating multiple parameters relevant for frame restoration to help gRNA evaluation and selection, as shown by gRNAs targeting *TMC1* c.229delA (**g**), *PCDH15* c.16delT (**h**) and *USH1C* c.369delA (**i**).

## Discussion

Here we present an *in vivo* genome editing strategy exhibiting unprecedented correction efficiency of frameshift mutations in postmitotic system, which is based on the highly biased, NHEJ-mediated editing outcome. In our *av3j* case, nearly 75% genome edited hair cells restore mechanotransduction, and this highly effective cell-level recovery further ameliorate auditory and vestibular symptoms in this USH1F animal model. As NHEJ is the major DSB repair pathway in almost all cell types, including both dividing and non-dividing cells^27^, in theory, this strategy enables efficient frame restoration in virtually any cells/organs. Besides, no editing template is required for this strategy, which is the necessity for HDR-mediated precise editing, thus it largely eases the strategy design and delivery procedure.

Our gRNA screening data reveal that gRNAs shifted in only 1-bp may exhibit totally different editing profiles with distinct dominant indels and indel size distributions, e.g. m-*3j*-gRNA1, m-*3j*-gRNA21 and m-*3j*-gRNA24 (Fig. 1b,c). Though some of these gRNAs may generate identical editing products, the final function-rescue fractions vary a lot for the different product frequencies, e.g. w-*USH1C*-369dA-g3 and w-*USH1C*-369dA-g4 (Fig. 6c,f,i). And in many cases, more than one gRNA are available for one frameshift mutation. These results demonstrate that the NHEJ-mediated frame-restoration strategy also enables highly flexible choice for therapeutic gRNAs.

In the past decade, great achievement has been obtained on gene therapy of inherited deafness. Gene replacement is the most common and successful type of genetic manipulation in this field^28–33^, which however is limited by the loading capacity of delivery vectors, e.g. 4.7 kb for AAV, rendering gene therapy for large proteins almost impossible. Recent progress on genome-editing-based *in vivo* deafness gene therapy showed significant hearing improvement on progressive hearing loss models, which harbor missense mutations on *TMC1* (cDNA 2.3kb), using either reading-frame disruption of dominant-negative allele^34,35^ or base editing^36^. Here, we provide the first study of large protein frame-restoration in a congenital profound deafness model, and also prove the possibility of USH1F amelioration by postnatal genetic manipulations.

Before our study, there are several studies also leveraged the non-random CRISPR cleavage and repairing patterns to correct pathogenic mutations^12,15^. MMEJ-mediated larger deletions were used to rescue pathogenic microduplication to wildtype sequences, which are relatively controllable for one could search for the microhomology arms and precisely predict the editing outcomes. Here, we take advantage of the NHEJ-mediated insertions and deletions, which are the dominant components of most editing outcomes, to compensate the small indels in frameshift mutations while leaving 1-2 amino acid alterations in the frame-restored products. Our data reveal that for some less important regions, imperfect correction could also mediate a significant phenotypic restoration.

During our explorations in *av3j* treatment, we find that for recessive mutations in diploid mammalian cells, only one functional allele is enough to achieve a cell-level functional rescue. Thus, gRNAs with 30% functional rescued products could provide about 50% function restoration cells containing diploid genomes. In our 114 gRNA screening, 55 of them could achieve this criterion, covering 65% (26/40) selected deafness mutations. If the percentage of function-restored cells are above the ‘minimal requirement’ for a system to conduct physiological function, a phenotypic restoration could be achieved. Once a gRNA could provide enough function-restored cell fraction for the targeted system, it deserves to try for *in vivo* applications.

Though we got some auditory function restoration in the USH1F mice, yet the rescue effect is still limited. Combining the nearly 75% functional recovery rates (Fig. 3b-d and Supplementary Fig. 4a,b) with the 100% and 70% transfection ratios in IHCs and OHCs (Supplementary Fig. 2f), around 75% IHCs and 53% OHCs could be recovered in the injected mice, thus the majority of hair cells were rescued by our treatment. However, unrecovered and disorganized hair bundles were observed along the entire cochleae in all tested, virus-delivered mice, which may also underlie the smaller mechanotransduction currents in targeted OHCs (Fig. 3c). And the irreversible cell death in middle and basal turn may further indicate that PCDH15 is not only important for hair-bundle development and organization, but also relevant for hair-cell maintenance and survival. Considering that the PCDH15 starts expression as early as embryonic day 12^37^, further study is needed to map the best therapeutic window for this protein.

Here we demonstrate the first *in vivo* gene therapy for frameshift mutation in a postmitotic system, and provide a practical guide for frame-restoration gRNA evaluation and selection. We anticipate that, with further improvement in profile prediction of mutant genomes and functional evaluation on edited products, this NHEJ-mediated frame-restoration strategy holds great promise for clinical treatment of small indels induced frameshift mutations, which consists 22% of the inherited Mendelian disorders in humans^38^.

## Methods

### Animals

All animal experiments were performed in compliance with the guidelines provided by the Institutional Animal Care and Use Committee (IACUC) at Tsinghua University. The mice were maintained in the Animal Research Facility in campus, which has been accredited by the Association for Assessment and Accreditation of Laboratory Animal Care International (AAALAC). Both *av3j/av3j* mice^18^ (C57BL/6J-*Pcdh15^av-3J^*/J, JAX:002072) and *v2j/v2j* mice^39^ (C57BL/6J-*Cdh23^v-2J^*/J, JAX:002552) were gifts from Dr. Ulrich Müller. The *av3j/av3j;Cas9+* mice were achieved by crossing *av3j/av3j* mice with the Cas9 knockin mice^22^ (Gt(ROSA)26Sor^*tm1.1(CAG-Cas9*-EGFP)Fezh*^, JAX:024858, Bar Harbor, ME). The mice were housed in a temperature-controlled room with 12-hr light/dark cycle and have free access to water and food. Both female and male mice were used.

### Design of gRNAs and plasmid construction

All gRNAs were constructed with the PX330 backbone (42230, Addgene). The primers used for plasmid construction were designed by CCtop (CRISPR/Cas9 target online predictor, https://crispr.cos.uni-heidelberg.de) with default parameters, except for setting the gRNA length to 17 nt. The plasmids were constructed as previously reported^40^. Briefly, the oligos were phosphorylated by T4 Polynucleotide Kinase (M0201S, NEB) at 37°C for 30 min, then annealed by temperature ramping from 95°C to 25°C in a thermocycler. The PX330 backbone was digested with BbsI (R0539S, NEB), and ligated with the oligos by T4 ligase (15224017, Invitrogen). The plasmids were extracted by MaxPure Plasmid HC kit (P1231, Magen, China) for following procedures including cell-line transfection and tissue electroporation.

### Cell/tissue culture and transfection

The N2a cell line was a gift from Dr. Yichang Jia (Tsinghua University), and was maintained in normal DMEM (C11965500BT, Thermo Fisher) containing 15% fetal bovine serum (AB-FBS-0500s, ABW) and 100 U/ml penicillin-streptomycin (15140148, Thermo Fisher). The N2a cells were seeded in 48-well plate in ~70% confluence at 24 hr before the transfection. The plasmids were transfected with Lipofectamine 2000 (11668019, Thermo Fisher) following manufacturer’s instructions. Normally, N1-EGFP was co-transfected to guarantee successful transfection.

Cochlear culture and injectoporation were as previously described^21^. In this study, the organ of Corti was dissected out from mouse pups and was attached on the inner side of a 35-mm culture dish lid cultured with 2ml DMEM/F12 medium (21041-025, Thermo Fisher) with ampicillin (1.5 μg/ml, GG101, TransGen Biotech) addition. For electroporation, plasmids (1 μg/μl in 1x HBSS) were injected into gaps between hair cells via a glass electrode with an open tip of 2 μm diameter. A series of 3 electrical pulses (80-V amplitude, 20-ms duration, 1-s interval) were immediately applied by an electroporator (BTX ECM830, Harvard Apparatus). After electroporation, the tissues were cultured in a humid incubator (37°C, 5% CO_2_).

### Immunofluorescence

For PCDH15 immunostaining, cultured or fresh dissected cochlear tissues were pre-treated with phosphate buffered saline (PBS, cc0005, Leagene; calcium free) adding 5 mM EGTA (E3889, Sigma-Aldrich) for 10 min at room temperature (RT), then fixed in 4% paraformaldehyde (PFA, DF0135, Leagene) for 10 min at RT. Then washed for 5 min x 3 times and blocked in 1% BSA (A3059, Sigma-Aldrich) (w/v) for 1 hour at RT before incubating with antibodies. Primary antibody: rabbit anti-PCDH15 (PB811, 1:400 dilution, a gift kindly provided by Dr. Ulrich Müller), incubate at 4°C for 16 hours followed by 15 min x 3 times wash in PBS. Secondary antibodies: goat anti-rabbit (A-21245, Alexa 647 conjugated, Thermo Fisher, 1:500) with either Alexa Fluor 488 (A12379, Thermo Fisher, 1:1000) or Alexa Fluor 568 Phalloidin (A12380, Thermo Fisher, 1:1000), incubate at RT for 2 hours followed by 15 min x 3 times wash in PBS. All antibodies were diluted in PBS with 5 mM EGTA by v/v. The tissues were finally mounted with ProLong Gold Antifade Mountant (P36930, Thermo Fisher). The images were acquired using a deconvolution microscope (DeltaVision, Applied Precision) equipped with a 100x oil-immersed objective and deconvolved using the Huygens deconvolution software (version 18.10, SVI).

For virus transfection and hair cell survival quantification, animals were sacrificed and harvested the inner ear tissues. Then fixed in 4% PFA for 30min at RT, washed in PBS by 7 min x 3 times. Dissected the basilar membrane and blocked in 1% FBS for 1 hour at RT before incubating with antibodies. Primary antibody: rabbit anti-Myo7a (25-6790, Proteus Biosciences Inc, 1:1000) incubate at 4°C overnight and followed by 15 min x 3 times wash in PBS. Secondary antibodies: goat anti-rabbit Alexa 647 conjugated at 1:2000 with Alexa Fluor 488 Phalloidin (1:1000). The tissues were finally mounted with ProLong Gold Antifade Mountant. The images were captured using Andor Dragonfly spinning disk confocal microscopy by the 40x oil-immersed objective. The spot and filament function of Imaris Software v9.3.1 (Oxford Instrument) were used for cell number quantification. For virus transfection quantification, gamma values were set to 3, in order to show clear labeling in the weak viral expression cells.

For functional tests on HEK293T cells, cells were transfected with EGFP-fused harmonin with or without mutations. Harvest 24hr-post-transfection, followed by 3 wash in PBS, and fixed in 4% paraformaldehyde (PFA, DF0135, Leagene) for 15 min at RT. Then washed 3 times in PBS. And stained with Phalloidin-Alexa Fluor 568 (A12380, Thermo Fisher, 1:1000) at 4°C overnight. Wash 3 times in PBS. Images acquired by Zeiss LSM780 confocal microscope using 100X (NA 1.4) oil-immersion objective. Maximum projections of z-stacks were handled using Zeiss Black software.

### Tissue analysis

#### Cochlea calcium imaging

The genetically encoded Ca2+ indicator GCaMP6m-X^41^ was co-transfected in cochlear hair cells. For calcium imaging, the tissues were bathed in fresh external solution (in mM): 144 NaCl, 5.8 KCl, 2.5 CaCl_2_, 0.9 MgCl_2_, 10 HEPES, 5.6 D-Glucose, PH7.4. A series of three fluid-jet stimuli (20 psi, 0.1 s, 0.3 s and 0.5 s) were applied by a glass capillary with an open tip of 5 μm diameter at 40-s interval. Images were acquired with an upright microscope with a 60x water-immersed objective (Olympus BX51WI) at 2-s sampling rate.

#### Whole-cell electrophysiology of hair cells

At assigned ages, the mice were sacrificed and their apical part of basilar membrane or whole saccules were taken out and transferred into dissection solution, which contains (in mM): 5 KCl, 140 NaCl, 1 MgCl_2_, 0.5 MgSO_4_, 0.1 CaCl_2_, 10 HEPES, 3.4 L-Glutamine, 10 D-Glucose, pH 7.2. The tectorial membrane or otolith was removed with fine forceps, then the organ of Corti or sacculus was transferred into a recording dish with recording solution, which contains (in mM): 145 NaCl, 0.7 NaH_2_PO_4_, 5 KCl, 1.3 CaCl_2_, 0.9 MgCl_2_, 10 HEPES, 5.6 D-Glucose, pH 7.4. Then mCherry expressed OHCs or VHCs were whole-cell recorded and membrane currents were measured by an electrophysiology amplifier (EPC-10 USB, HEKA, Germany) at sampling rate of 20 kHz. The cells were held at –70 mV and the hair bundle was stimulated by a fluid jet. For OHCs, 40-Hz sinusoid stimulation in 60 ms was performed. For VHCs, a 500-ms square wave was used in order to obtain large enough stimulation.

#### FM5-95 loading and imaging

Wild type C57B6J animals and *av3j/av3j;Cas9+* animals with or without virus injection were sacrificed at P7-P10. Basilar membranes were dissected in 1 X HBSS and bathed in 6uM ice-cold FM5-95 (T23360, Thermo Fisher, dissolved in 1 X HBSS) for 30s, then followed by 4-5 washes in clean HBSS. Tissues were attached to glass bottom dish upside down by prolong-antifade solution. Images were obtained using Zeiss LSM880 confocal microscope equipped with standard laser lines (405, 514, 561, 633 nm) using 40 X (NA 1.2) water-immersion objective. Maximum projections of z-stacks were handled using Zeiss Black software. FM5-95 was excited by 514-nm laser and monitored in the region between 680-742nm, an emission wavelength to distinguish from GFP signal (Cas9 KI) and mCherry signal from virus.

### Genome editing analysis

#### Tissue digestion and FACS

Electroporated or AAV-transfected auditory tissues (containing basilar membrane, spiral ligament, and stria vascularis) and vestibular tissues (include utricles, saccules and semicircular canals) were dissected in ice-cold HBSS (C14175500CP, Thermo Fisher) quickly and then incubated in HBSS with 1 mg/ml collagenase I (C0130-1g, Sigma-Aldrich) and 0.05% trypsin-EDTA (25200056, Thermo Fisher) at 34°C for 10-15 min. The effect of enzymatic dissociation was checked with an inverted microscope. Then equal volume of 10% FBS-containing DMEM/F-12 medium was added to stop the reaction. After centrifuging at 1,000 g for 1min, the cells were washed with 10% FBS-containing DMEM/F-12 and re-suspended. After passing through a 70-μm filter to remove large debris, the isolated cells with fluorescence signals were sorted and enriched directly into 0.2-μl PCR tubes by a FACS machine (BD Influx). Noted that, in order to get enough cells for downstream deep sequencing, the FACS gate was relatively loose to collect total cells containing red cells at 5-20%.

#### Deep sequencing library construction and data analysis

Genome DNA of transfected N2a cells or tissue cells were extracted by QuickExtract™ DNA Extraction Solution 1.0 (QE09050, Lucigen, Middleton, WI) following the manufacturer’s instructions. For each targeting genome locus, an amplicon of 100-300 bp was designed, with cleavage sites located about 50-100bp away from amplicon boundaries. Firstly, the first run of PCR (PCR1) was applied to amplify targeting DNA with primers containing part of the Ilumina adaptors. And exonuclease I was added to the PCR1 products to clear all remaining PCR1 primers. Then a second run of PCR was performed to add unique barcodes and P5/P7 flow cell binding sites. All the PCR reactions were carried out with Kapa HiFi 2 x mastermix (KK2601, Kapa Biosystems). PCR products were then separated by 2% agarose gel, and only products with desired sizes were cut and purified by Gel Purification kit (D2110, Magen, China). After quantified by Qubit™ 1X dsDNA HS Assay Kit (Q33230, Thermo Fisher) and verified by Agilent 2100 Bioanalyzer, samples were pooled together and sequenced 2X150 on the Illumina HiSeq X-Ten platform. Sequencing reads were demultiplexed according to the 8-nt index sequence and then individual FASTQ files were analyzed by the batch version of CRISPResso2 v.2.0.32^42^ with the CRISPRessoPooled mode. The analysis parameters were set as: -w 12, -q 30, -s 0, --max_paired_end_reads_overlap 150, while others were used as default. And all other analysis and quantification were based on the “Alleles_frequency_table.txt” in the CRISPResso2 report with custom Python scripts or Excel2016. Noted that, all our editing analysis exclude reads with only substitution. The indel frequency were calculated as (all edited reads – substitution only reads)/(all reads for the target locus).

#### Off-target prediction

The mismatch off-targets were predicted by CCtop^43^ (CRISPR/Cas9 target online predictor, https://crispr.cos.uni-heidelberg.de). All the mismatched off targets were ranked by number of mismatches, then the off-target loci were chose with different alignment parameters. The off targets with DNA bulges were predicted by Cas-OFFinder^44^ (http://www.rgenome.net/cas-offinder/), set the DNA bulge size to 1bp. Off-target loci with different bulge locations were chosen. Noted that, all selected bulge loci also had 1 bp mismatch with the on-target site. And all off-targets avoided repeating region.

#### Function prediction of editing product

We used “Alleles_frequency_table.txt” in the CRISPResso2 report as the input files for the function prediction analysis. The “MODIFIED” reads were first aligned with WT CDS sequences to obtain the CDS region. Then create the predicted editing products in mutant genomes by manually adding the ±1-bp mutations into the editing reads based on WT genomes using following rules: for 1-bp deletion mutations, if the deletion positions were not edited, delete the base at the deletion position, otherwise, delete the bases closest to the 3-bp upstream the PAM sequences; for 1-bp insertion mutations, if the insertion positions were not edited, insert the bases in the insertion sites, otherwise inserted the mutant bases at positions that make the edited indels have closest distances with the 3-bp upstream the PAM sequences. Only the predicted products that were multiples of 3 would be translated into protein sequences and continued function prediction.

Each amino acid position was assigned to the conservation score (rank from 1 to 9, from the highly variable to highly conserved, corresponded to the 9 color codes in figures) calculated by ConSurf 2010^45^ (http://consurf.tau.ac.il) using parameters: multiple sequence alignment using MAFFT; homologues collected from UNIREF90; homology search algorithm using HMMER (E-value 0.0001, No. of iteration 1); maximal %ID between sequences:95; minimal %ID for homologs:35; using 150 sequences that sample the list of homologues; method of calculation: Bayesian; model of substitution for proteins: best fit.

And the predicted product would be predicted as nonfunctional if it satisfy any of following criteria:

1. Containing any stop codons;
2. For amino acid deletion: deleted highly conserved amino acids (conservation score ≥7) and or deletion happen near highly conserved amino acids (one or both neighboring amino acid score =9);
3. For amino acid insertion: insertion happen in between highly conserved amino acids (both neighboring amino acid score ≥7);
4. Total amino acid changes ≥ 3, including substitutions, insertions and deletions;
5. Substitution of highly conserved amino acids (conservation score ≥7), and the amino acid switches did not exist during evolution;
6. Substitution of highly conserved amino acids (conservation score ≥7), and the amino acid switches change the acid-base property of the amino acid position.

#### Specificity prediction of gRNAs

The gRNA specificity were calculated in Benchling (https://www.benchling.com/), which enable scoring gRNA with 17-nt length. A uniform formula, based on off-target mismatch and mismatch position^46^, were used to calculate a specificity score for each gRNA, thus enabled a comparison of gRNA specificity even in different loci.

#### Radar plot drawing

In the radar plot, the value of each axis was in linear distribution, with the max value located in the outermost ring. The maximal values of %Inframe, %Predicted aa change ≤ 3, %Predicted functional rescue were set to 100%. And the center was set to 0%. The value of the outermost ring was 15% for %Indel and 45 for specificity.

### Animal treatment and analysis

#### Surgery and virus injection

All AAVs used in this study were customized from Vigene Bioscience, Jinan, China. For anesthesia, P0-1 mice were put on ice and waited for 1-2 min. Then dorsal incision was made, and the ear was approached to expose the bulla. With a stereotaxic microscope, the scala media was visually located between the round widow niche and the stapedial artery, then 600-nl virus solution (1.5 × 10^14^ vg/ml) was injected into scala media at a rate of 250 nl/min. Fast green (F7258, Sigma-Aldrich) was added to visualize delivery effect, as successful delivery to scala media came with a ribbon pattern. For each pup, both ears were injected and all procedures were finished within 20 min.

#### Auditory brainstem response (ABR)

Mice were anesthetized by intraperitoneal injection of tribromoethanol (300 mg/kg). The recording electrode was inserted underneath the scalp right between the two ears. The reference electrode and the ground electrode were inserted subcutaneously at the pinna and the groin respectively. ABR data were collected by an RZ6 workstation controlled by a BioSig software (Tucker-Davis Technologies, Alachua, FL). Pure tone stimuli was applied at 4, 5.6, 8, 11.2, 16, 32 kHz. Maximum stimulus intensity was set to 90 dB SPL, with attenuation decreasing from 110 dB to 10 dB SPL at 5-10dB steps.

#### Vestibulo-ocular reflex (VOR) and data analysis

The VOR recording and data analysis was conducted as previously reported^47^. Briefly, in a metal chamber, the mouse was immobilized by a clamp and mounted on a mirror-camera-equipped turn table. Then the binocular eye movements were evoked by ± 20° sinusoidal rotations at 0.25, 0.5, or 1 Hz frequencies. For each animal, a 2-3 min video was recorded and traced by a custom machine-learning-based scripts in matlab R2016a. The horizontal eye-move degree was calculated as

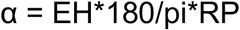

Where α indicates horizontal eye-movement degree, EH indicates horizontal eye-movement distance, and pi means the Constant of PI, and RP indicates the axial length of eye-movement^48^. Six periods, whose peak frequency coincided with stimulus frequency, were averaged to calculate the final eye movement angles. The gain value was calculated as:

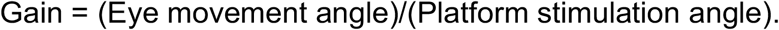

#### Open field test and data quantification

4~8 week-old animals were used for all behavior tests. A 33 × 33 cm arena was used with infrared illumination. Animals were placed on one side of the arena and videotaped for 10 min. The arena was cleaned before changing to another animal. The circling behaviors were quantified by a behavior analysis software (The EthoVision XT version 11.5, Noldus Information Technology). Both clockwise and anticlockwise circling were counted in analysis of circling frequency. And the moving traces are tracked by custom MATLAB scripts.

#### Rotarod test

Before test, animals were trained on a rotarod (47650, Ugo Basile) for 5 times in 2 days, by which first two workouts used constant 10 r.p.m for XX s, and the next three were the final 300-s test program that began at 10 r.p.m and then increased to 30 r.p.m in 90 s. During test, each animal ran 5 times to count the running time before it fell from the rotarod. After excluding the maximum and minimum, the averaged running time was used for data presentation.

#### Acoustic startle response

The acoustic startle responses were recorded by Xeye Startle Reflex system (v1.2, Beijing MacroAmbition S&T Development). The animals were tested in a sound shielded startle box and the startle responses were sensed by the gravitational acceleration sensor fixed beneath an elevated platform. Mice were placed in a smaller, squared, plastic chambers and anchored to the sensing platform during recording. Experiment started with a 5 min 60dB background white noise, and then followed by 32 trials of single noise pulse presented at pseudorandom order at randomized inter trial intervals between 10-50s (90 to 120dB with 10 dB steps, 40ms duration with 0ms onset and offset ramps). The fan was opened during whole experiment to reduce external noise interference. Eight repetitions were recorded for each sound intensities per each subject.

### Data analysis

Charts were plotted by Prism 6 (GraphPad Software, San Diego, CA). All data are mean ± SEM or SD, as indicated in the figure legends. ANOVA tests were used to determine statistical significance (*\*P* < 0.05, *\*\*P* < 0.01, *\*\*\*P* < 0.001, *\*\*\*\*P* < 0.0001).

## Acknowledgements

We thank Yanhui Chen and Dr. Bai Du in device usage for acoustic startle response and rotarod, thank Dr Yinqing Li, Xin Liang, Chunlai Chen, and members of Xiong laboratory for helpful discussions and critical proof-reading of this manuscript, thank the Imaging Core Facility, Technology Center for Protein Sciences at Tsinghua University for assistance of using imaging instruments and software, thank the Tsinghua University Branch of China National Center for Protein Sciences (Beijing) and Tsinghua University Technology Center for Protein Research for the cell function analyzing facility support, thank Core Facility of Center of Biomedical Analysis, Tsinghua University for assistance with L780 confocal microscopy.

This work was supported by the National Natural Science Foundation of China (31522025, 31571080, 81873703, and 3181101148), Beijing Municipal Science and Technology Commission (Z181100001518001), and a startup fund from the Tsinghua-Peking Center for Life Sciences.

## Author contributions

L.L. did gRNA design, construction, FACS and deep sequencing, tissue culture and injectoporation, mouse surgery and viral injection, behavioral test; K.L. did genome data mining; L-Z.Z. performed the cochlear culture and injectoporation, immunostaining and plasmid preparation; H-Q.H. did behavioral test and immunostaining data quantification; Q.H. performed the cochlear culture and injectoporation; S.L. did the hair-cell electrophysiology and behavioral data analysis; S-F.W., Y-Z.W., J.L., C-M.S., J-F.C., C-R.L., H-B.D., and J.L. provided necessary help; F-Y.C., Z-G.X., W-Z.S. and Q-W.S. supervised some of the experiments; L.L. and W.X. designed experiments, generated figures, and wrote the manuscript; W.X. supervised the project.

## Notes

### Competing Interest Statement

The authors have declared no competing interest.

